# Regulation of the aurantio-obtusin accumulation by StTCP4.1-mediated StDA1-StHDR1 module in *Senna tora* seeds

**DOI:** 10.1101/2024.01.08.574662

**Authors:** Shuang Liu, Jinling Liu, Ann Abozeid, Xuecui Yin, Juane Dong, Zongsuo Liang

## Abstract

*Senna tora* (*S. tora*) is a commonly used Chinese medicinal plant due to the presence of the bioactive compounds anthraquinones in its mature seeds. Seed size is an important factor that affect *S. tora* yield quantity and quality. However, the mechanism regulating seed size and aurantio-obtusin biosynthesis in *S.tora* is still unclear. In this study, we identified the ubiquitin receptor StDA1 in *S.tora* that served as a negative regulator to seed formation and limited seed enlargement. Antisense overexpression of StDA1 led to larger seeds in *S. tora* and promoted the accumulation of aurantio-obtusin. In contrast, overexpression of StDA1 in *S.tora* resulted in a relative decrease in aurantio-obtusin accumulation. Moreover, StDA1 can directly bind to StHDR1and regulate its degradation through the 26S proteasome to regulate seed size and aurantio-obtusin accumulation. We also found that the StDA1-StHDR1 module is responsive to the MeJA via StTCP4.1, which in turn affects the accumulation of aurantio-obtusin. Overall, we have identified a protein complex that regulates the accumulation of aurantio-obtusin, StTCP4.1-StDA1-StHDR1, as a potential target for improving *S.tora* yield quantity and quality.

## Introduction

*Senna tora* (*S.tora*) is a commonly used Chinese medicinal plant that has a high medicinal and nutritional value. In food industry, *S. tora* seeds were adopted as an ingredient for the production of beverages and teas(Wan et al, 2016); in the medical field, *S.tora* seeds were applied in the treatment of numerous diseases including cardiovascular diseases, psoriasis, diabetes, and dermatological disorders(Guo et al, 2017; Ko et al, 2021; Meena et al, 2010). This pharmaceutical action is owing to the presence of anthraquinones in mature seeds(Fernand et al, 2008; Kang et al, 2020). However, the qualification rate of anthraquinone content in *S. tora* seeds in the current market was reported to be only 33.3%(Bao et al, 2018). So, with the growing demand for *S.tora* seeds in food and pharmaceutical industries, there is an urgent need to increase the yield quantity and quality

Seed size is not only an important indicator to measure the seed yield of crops but also a key trait affecting seed dispersal and seedling establishment(Gegas et al, 2010; Kesavan et al, 2013; Zhang et al, 2015). The coordinated growth of the seed coat, endosperm, and embryo is regulated by both maternal and zygotic tissues, which establish the seed size(Li et al, 2019). In *Arabidopsis thaliana* (*A. thaliana*), plant hormones and the HAIKU (IKU) pathway regulate seed size by modulating the growth of zygotic tissues(Gehring & Satyaki, 2017). For example, premature endosperm cellization in the *iku* mutant results in smaller seeds(Luo et al, 2005; Wang et al, 2010), while the regulation of seed size by the phytohormones ABA and BR are mediated through the IKU pathway(Cheng et al, 2014; Jiang et al, 2013).

In addition, the ubiquitin-proteasome pathway was found to be one of the major signaling pathways controlling seed size by maternal tissues. The pathway consists of ubiquitin-activating enzyme (E1), ubiquitin-coupled enzyme (E2), and ubiquitin ligase (E3)(Shu & Yang, 2017), and the E3 ligase is used for specific recognition and combining with substrates(Liu et al, 2021). In *A. thaliana*, DA1 is recognized as a ubiquitin receptor with two ubiquitin interaction motif (UIM) structural domains and a LIM structural domain at its N-terminus, as well as a peptidase structural domain at the C-terminus(Dong et al, 2017; Li et al, 2008). The modulation of cell proliferation by the DA1 pathway is crucial in the regulation of seed size in the maternal system. DA1 and its homolog da1-related gene (DAR1) can negatively regulate seed size. The simultaneous deletion of DA1 and DAR1 leads to larger seeds. The E3 ligases DA2 and ENHANCER OF DA1 (EOD1)/BIG BROTHER (BB), as well as the deubiquitinating enzyme UBIQUITIN-SPECIFIC PROTEASE15 (UBP15)/SUPPRESSOR2 OF DA1 (SOD2) are all able to interact with DA1 to synergistically regulate seed and organs size(Dong et al, 2017; Du et al, 2014; Xia et al, 2013). Genetic analyses have shown that UBP15 has opposite roles to DA2 and EOD1/BB in regulating seed and organs size(Li et al, 2019).

In other species, the DA1 pathway also plays an essential role in the regulation of seed size. For instance, down-regulation of BnDA1 in *Brassica napus* increased its seeds yield (Wang et al, 2017); overexpression of mutated ZmDA1 or ZmDAR1 in maize increased grains yield(Xie et al, 2018); and similarly, overexpression of TaDA1 reduces the size and weight of wheat grains(Liu et al, 2020). However, in rice (*Oryza sativa*), the homolog of AtDA1, the ubiquitin-interacting motif-type ubiquitin receptor (HDR3) positively regulates grain size by promoting spikelet shell cell division(Gao et al, 2021). In conclusion, since the DA1 pathway enhances seeds yield, it is important to characterize the DA1 pathway and other components in a variety of species and to measure their contributions to crops yield.

Jasmonic acid (JA) is a lipid (oxylipin) plant hormone that is involved in the regulation of plant development, secondary metabolites accumulation, abiotic stress response, fruit ripening, and other processes. The TEOSINTE BRANCHED 1 / CYCLOIDEA / PROLIFERATING ELL FACTOR (TCP) transcription factor (TF) plays a critical regulatory function in the biosynthesis of JA and other oxylipins. Statistics show that 8 of the 19 oxylipin synthesis genes (including LOX2) contain a TCP binding site (TBS) on the promoter(Schommer et al, 2008). It has been found that AtTCP4 and AtTCP20 can combine with the LOX2 promoter to enhance the expression of this gene(Danisman et al, 2012), which promotes leaf senescence in *A. thaliana*. In *Artemisia annua*(*A. annua*), AaTCP14 and AaTCP15 are involved in the regulation of the JA signaling pathway, and artemisinin content was significantly elevated in AaTCP14 and AaTCP15 overexpressed plants, while it was significantly reduced in silenced plants(Ma et al, 2018; Ma et al, 2021). Furthermore, in *S.tora*, StTCP4.1 TF was also identified in response to the JA signaling pathway. StTCP4.1 responds to JA signaling mainly by interacting with StMYC2a(Liu et al, 2022). However, the mechanism of StTCP4.1 in regulating the formation of *S.tora* seeds and aurantio-obtusin accumulation has not yet been reported.

In our study, we identified an AtDA1 ortholog, StDA1, whose expression was significantly down-regulated in the late stage of seed maturation. Therefore, we hypothesized that StDA1 may be involved in seed formation and aurantio-obtusin biosynthesis. Firstly, we characterized the ubiquitin receptor gene StDA1 from *S.tora*, and found that StDA1 inhibited plant growth and led to smaller seeds, and reduced the accumulation of aurantio-obtusin. Secondly, we revealed that StDA1 restricts the biosynthesis of aurantio-obtusin by regulating the protein stability of StHDR1, a key enzyme in the aurantio-obtusin biosynthesis pathway. Additionally, the negative regulation of seed formation and aurantio-obtusin biosynthesis by the StDA1-StHDR1 module depends on the intervention of MeJA-mediated StTCP4.1, which is able to inhibit the facilitating effect of StTCP4.1 on StDA1-StHDR1 interactions, thereby reducing the accumulation of aurantio-obtusin. In conclusion, the MeJA-mediated StDA1-StHDR1 regulatory module offers a deeper understanding to the molecular events underlying the formation of *S.tora* seeds and the biosynthesis of aurantio-obtusin.

## Results

### Identification and expression analysis of StDA1

DA1 was previously reported to regulate seed size in *A. thaliana*, maize, rapeseed, and wheat(Chen et al, 2021). To expand our understanding of DA1 in *S.tora*, we identified three genes of the DA1 family from the *S.tora* genome(Kang et al, 2020) by the BLASTP search based on the *A. thaliana* DA1 protein sequence(Li et al, 2008) as a query. We found that only one gene was a homolog of AtDA1, and the other two genes were homologs of AtDAR2 (Figure 1A). We named the DA1 homolog in *S.tora* as StDA1. The DA1 family is strongly conserved in plants (Figure S1 and Table S1). Protein structural domain analysis demonstrated that StDA1 contains two tandemly arranged ubiquitin interaction motif (UIM) domains at its N-terminus, a Lin11/ is-1/Mec-3 (LIM) domain located in the intermediate region, and a DA1-like structural domain at the C-terminus.

**Fig. 1.**
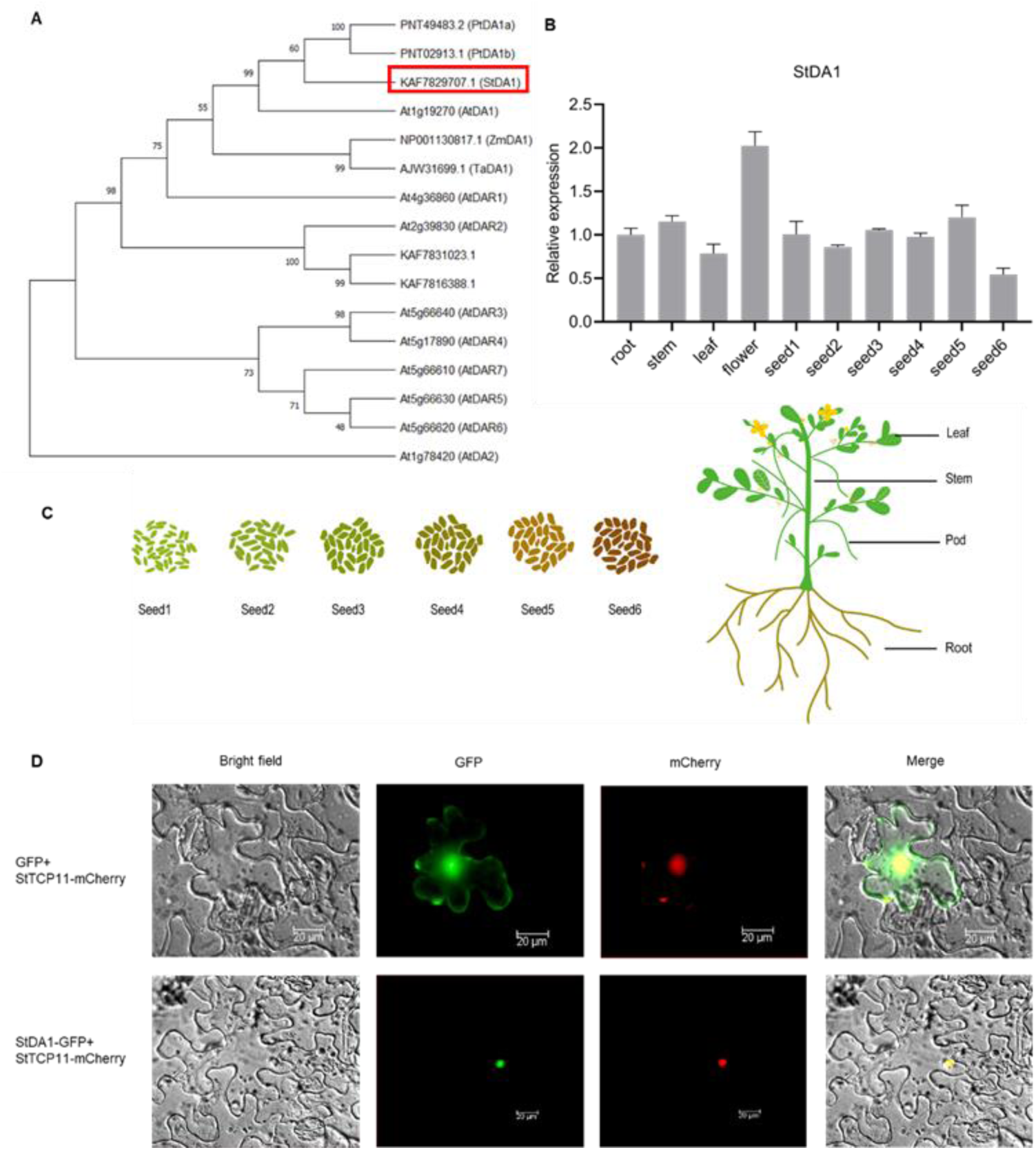
The StDA1 family and expression patterns of StDA1 in *S.tora*. (**A**) The evolutionary tree of the DA1 gene family from *Senna tora* (St), *Zea mays* (Zm), *Arabidopsis thaliana* (At), *Populus trichocarpa* (Pt), and *Triticum aestivum* (Ta). (**B**) RT-qPCR analysis of the StDA1 gene expression patterns in various tissues of *S.tora*. Bars are means ±SD. (**C**) The model diagram of different growth and development periods of *S.tora* plant and seed. (**D**) The subcellular location of StHDR1 in tobacco leaves. StTCP11 was cloned into pCAMBIA1300-mCherry as a control. Images were observed under a fluorescent inverted microscope. Scale bars = 20 μm.

To elucidate the expression pattern of the StDA1 gene in *S.tora*, we used the reverse transcription quantitative PCR (RT-qPCR) method to measure the relative expression of StDA1 gene in different tissues of *S.tora*. StDA1 expression level was highest in flowers and lowest in seeds at the late maturation phase (Figure 1B and C). Since StDA1 is a negative regulator and the aurantio-obtusin that accumulates in *S.tora* only in the seeds (Figure S2), we can predict that StDA1 may have a positive regulatory effect on the biosynthesis of aurantio-obtusin in *S.tora* seeds. To characterize the subcellular localization of StDA1 protein, we successfully cloned and ligated the full-length coding sequence (CDS) of StDA1 into the pCAMBIA1300-GFP vector to fuse it with green fluorescent protein (GFP). Driven by the 35S promoter, StDA1 was fused to GFP and expressed in tobacco leaves. The results displayed that StDA1 fused with GFP protein was expressed only in the nucleus (Figure 1D), indicating that StDA1 exerts an influential effect on the nucleus.

### StDA1 negatively regulates seed size and aurantio-obtusin accumulation

To further investigate the role of StDA1 in *S.tora* seed formation and accumulation of aurantio-obtusin, we overexpressed and antisense overexpressed StDA1 in *S.tora* under the control of the CAMV 35S promoter. We obtained three lines from StDA1OE and StDA1AE transgenic *S.tora* plants and also obtained transgenic *S. tora* plants with empty vector (EV) as control (Figure S3). Subsequently, we analyzed the agronomic phenotypic traits of StDA1OE, StDA1AE and EV plants. The plant height and plant width of StDA1OE lines were significantly lower than those of EV lines, whereas the difference in plant height between StDA1AE and EV lines was smaller, and the plant width of StDA1AE was significantly larger than that of EV lines (Figure 2A, D and E). At the pod-bulging stage of *S.tora*, we found no significant difference in the number of tillers in StDA1OE, EV, and StDA1AE lines (Figure 2F). The number of pods and the length of pods were slightly lower in StDA1OE than in the EV strain, whereas the results were the opposite in the StDA1AE lines (Figure 2B, G and H). At the maturity stage of *S.tora* seeds, the hundred-seed weight of StDA1OE lines was slightly lower than that of the EV lines, and the length and width of the seeds of the StDA1OE lines were not markedly different from those of the EV lines. While the hundred-seed weight and length of the seeds of the StDA1AE lines were slightly higher than those of the EV lines, the width of the seeds of the StDA1AE lines was not markedly different from that of the EV lines (Figure 2C, I, J and K). In a nutshell, StDA1 negatively regulates the growth of *S.tora* plants and seed formation.

**Fig. 2.**
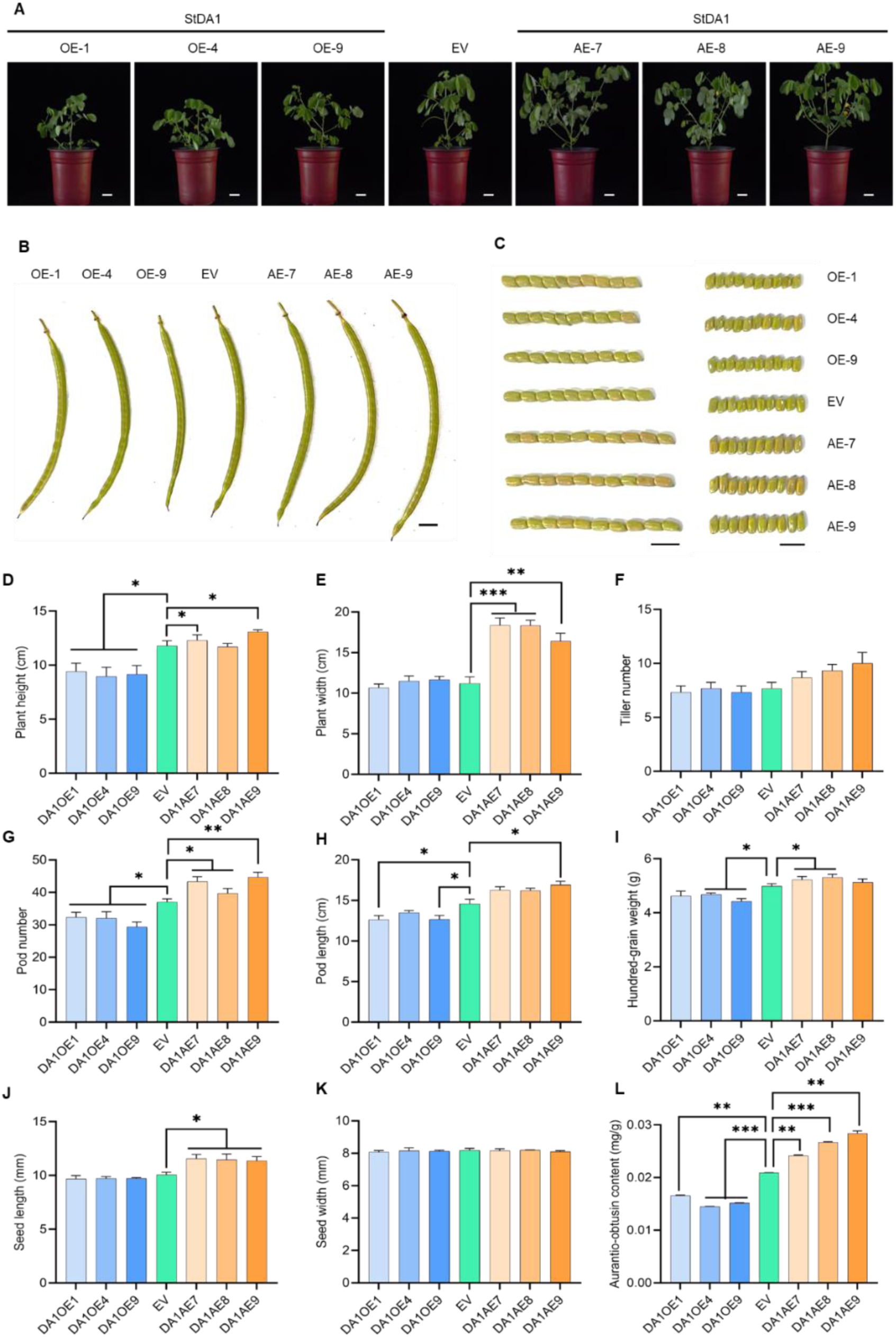
Overexpression and antisense overexpression of StDA1 affect plant growth in *S.tora*. (**A**) The plant phenotypic analysis of overexpression and antisense overexpression of StHDR1 lines, scale bars = 5cm. (**B and C**) The analysis of pod and seed size of overexpression and antisense overexpression of StHDR1 lines, scale bars=1 cm. (**D–L**) (**D**) the plant height; (**E**) the plant weight; (**F**) the tiller number; (**G**) the pod number; (**H**) the pod length; (**I**) the hundred-grain weight; (**J**) the seed length; (**K**) the seed width; and (**L**) the aurantio-obtusin content of overexpression, antisense overexpression of StDA1, and EV plant lines. Bars are means ± SD. Statistical analysis was carried out using one-way ANOVA with multiple comparisons, and significant differences were uncovered compared to EV (*, P < 0.05; **, P < 0.01, ***, P < 0.001).

Aurantio-obtusin is the main active ingredient in *S.tora* seeds, and it is only present in the seeds. To explore whether StDA1 affects the accumulation of aurantio-obtusin in *S.tora* seeds or not. Seeds of StDA1OE, EV and StDA1AE lines were collected at seed maturity stage and dried using an oven at 50 ℃. Subsequently, high-performance liquid chromatography (HPLC) experiments were conducted to detect the content of aurantio-obtusin in StDA1OE, EV, and StDA1AE lines. HPLC results showed a significant decrease in aurantio-obtusin content in StDA1OE lines and a significant increase in its content in StDA1AE lines compared to EV lines (Figure 2L), indicating that StDA1 negatively regulated the accumulation of aurantio-obtusin in *S.tora* seeds. In order to study the regulation of StDA1 on the biosynthesis pathway genes of aurantio-obtusin, we used RT-qPCR to detect the changes of these synthase genes at the transcriptional level in StDA1 transgenic plants. The results showed that the expression of synthase genes in shikimic acid pathway and polyketide pathway was irregular in StDA1 transgenic plants, while the expression of ACCA, HMGR1 and MK1 genes in MVA pathway was significantly increased in StDA1OE plants and significantly decreased in StDA1AE plants. The expression of DXS4 and HDR1 in MEP pathway were significantly decreased in StDA1OE plants and significantly increased in StDA1AE plants (Figure 3). Overall, the changes in the expression levels of DXS4 and HDR1 synthetase genes in the MEP pathway were in contrast to the changes in the content of aurantio-obtusin, indicating that StDA1 inhibited the accumulation of aurantio-obtusin may be related to DXS4 and HDR1.

**Fig. 3.**
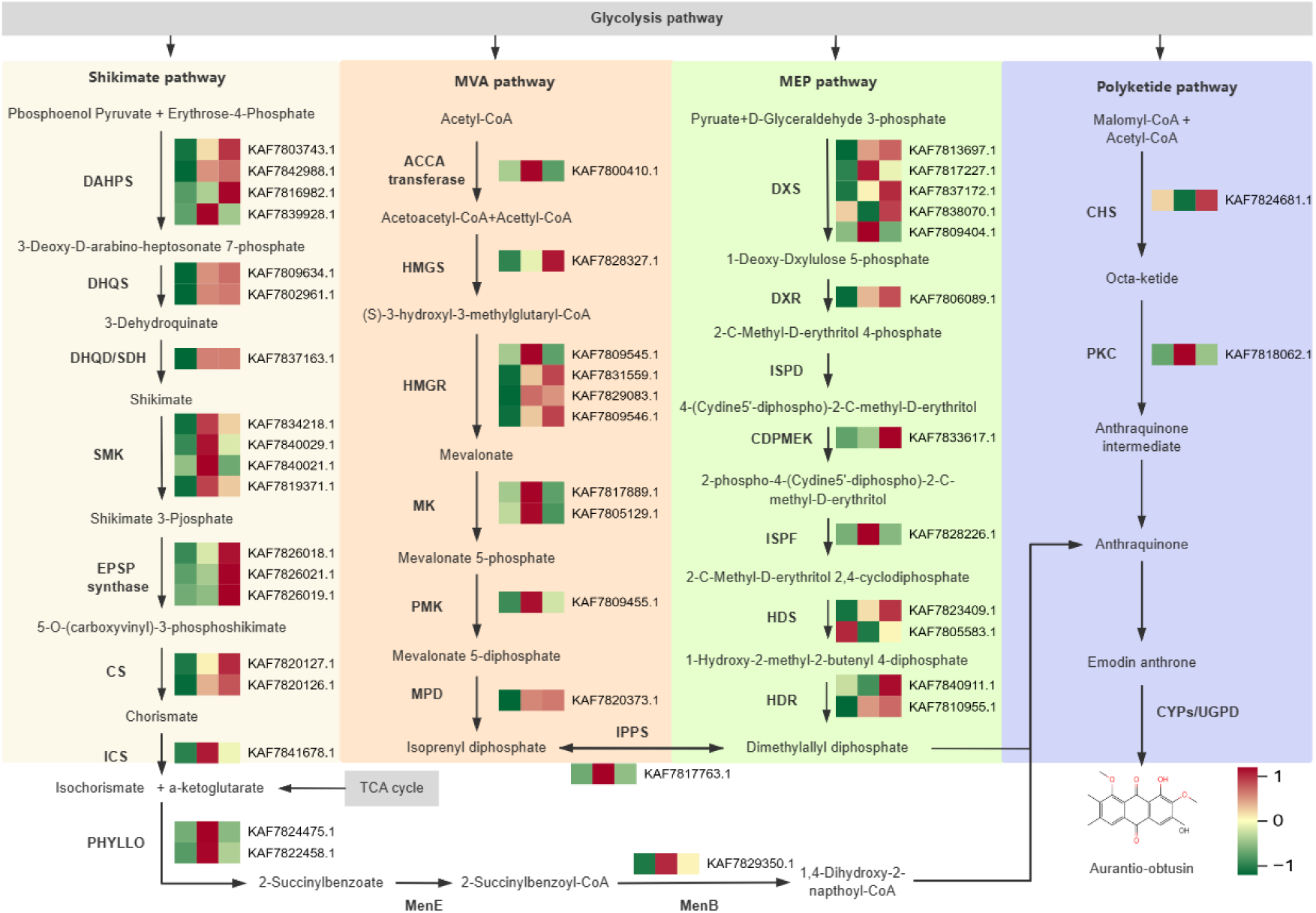
The expression levels of related genes involved in aurantio-obtusin biosynthesis in StDA1 transgenic plants. The heat map from left to right is empty vector, StDA1OE, and StDA1AE lines. The expression level of the gene in the empty vector plant was used as a reference. Red indicates that the gene expression is high, and green indicates that the gene expression is low. DAHPS, 3-Deoxy-7-phosphoheptulonate synthase; DHQS, 3-Dehydroquinate synthase; DHQD/SDH, 3-Dehydroquinate dehydratase/shikimate dehydrogenase; SMK, Shikimate kinase; EPSP, 3-Phosphoshikimate 1-carboxyvinyltransferase; CS, Chorismate synthase; ICS, Isochorismate synthase PHYLLO, 2-Succinyl-5-enolpyruvyl-6-hydroxy-3-cyclohexene-1-carboxylic acid synthase; MenE, 2-Succinylbenzoate-CoA ligase; MenB, 1,4-Dihydroxy-2-naphthoyl-CoA synthase; ACCA, Acetyl-CoA carboxylase; HMGS, Hydroxymethylglutaryl-CoA synthase; HMGR, Hydroxymethylglutaryl-CoA reductase; MK, Mevalonatekinase; PMK, Phosphomevalonate kinase; MPD, Methyl parathion hydrolase; DXS, 1-Deoxy-D-xylulose-5-phosphate synthase; DXR, 1-Deoxy-D-xylulose-5-phosphate reductoisomerase; ISPD, 2-C-Methyl-D-erythritol 4-phosphate cytidylyltransferase; CDPMEK, 4-Diphosphocytidyl-2-C-methyl-D-erythritol kinase; ISPF, 2-C-Methyl-D-erythritol 2,4-cyclodiphosphate Synthase; HDS, (E)-4-Hydroxy-3-methylbut-2-enyl-diphosphate synthase HDR, 4-Hydroxy-3-methylbut-2-enyl diphosphate reductase; IPPS, Isopentenyl-diphosphate delta-isomerase; CHS, Chalcone synthase; PKC, Polyketide cyclase/dehydratase.

### StDA1 can interact with the key enzyme StHDR1

To further explore the molecular mechanism underlying regulation of aurantio-obtusin accumulation in *S.tora* seeds by StDA1, we investigated its interaction partners using LC-MS/MS. As a consequence, the key enzymes StHDR1(4-hydroxy-3-methylbut-2-enyl diphosphate reductase) and StHDR2 were identified (Figure S4 and Table S2). Therefore, we performed Y2H experiments to examine whether StHDR1/2 could physically interact with StDA1. The Y2H experiments revealed that yeast cells co-expressing StDA1-BD and StHDR1-AD could grow on SD/-Leu-Trp-His-Ade medium containg 200 ng/ml Aureobasidin A (AbA), whereas StHDR2-AD could not (Figure 4A), indicating that only StHDR1 could interact with StDA1. To further validate these results, we performed additional BiFC, pull down and LCI experiments. BiFC analysis showed significant YFP fluorescence on tobacco leaves with StDA1-nYFP/StHDR1-cYFP co-expression, while YFP fluorescence was absent in the negative control (Figure 4B). Consistent with the BiFc assay, LCI assays were conducted to further confirm higher LUC activity and stronger luminescence signal in the StDA1-Cluc/StHDR1-Nluc co-expressed tobacco leaves but not in the negative control (Figure 4C-E), demonstrating that StDA1 can interact with StHDR1 in vivo. We then performed pull-down experiments with purified recombinant GST-tagged StDA1 (StDA1-GST) and His-tagged StHDR1 (StHDR1-His), and found that in addition to His-tagging, StHDR1-His proteins could pull down StDA1-GST, suggesting that StDA1 physically interacts in vitro with StHDR1 (Figure 4F). Taken together, StHDR1, a vital enzyme in the aurantio-obtusin biosynthesis pathway in *S.tora*, can interact with StDA1, suggesting that StDA1 may regulate the accumulation of aurantio-obtusin in *S.tora* seeds through the key enzyme StHDR1.

**Fig. 4.**
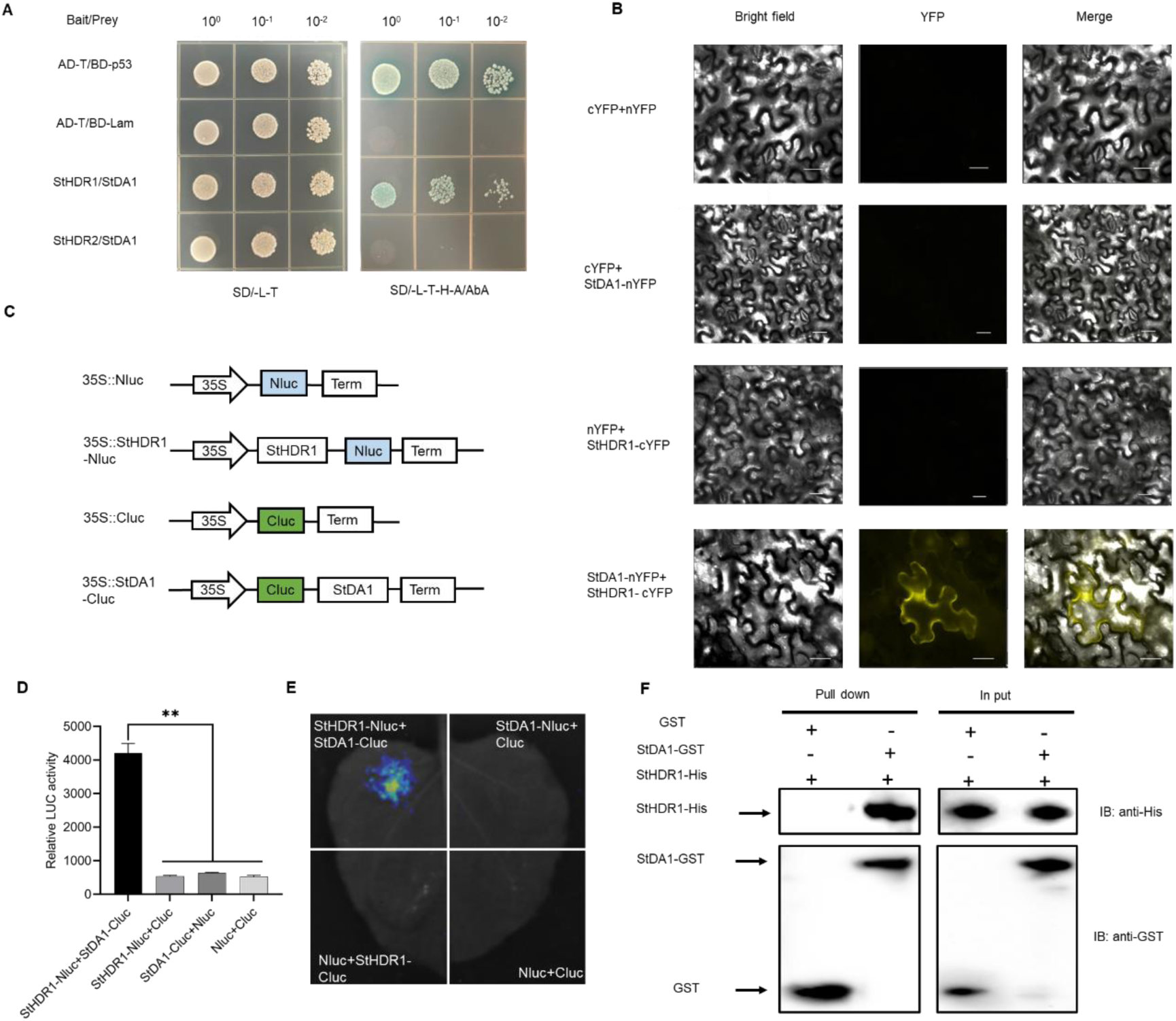
StDA1 directly interacts with StHDR1. (**A**)The interaction of StDA1 with StHDR1/2 in Y2H assays. The yeast cells were observed after 4 days of growth on SD/-Trp-Leu and SD/-Trp-Leu-His-Ade medium containing AbA and X-a-gal. (**B**) The interaction of StDA1 with StHDR1 in BiFC assays. Fluorescent signals were showed in tobacco leaves co-expressed with StDA1-nYFP and StHDR1-cYFP. Co-expression of StDA1-nYFP/cYFP, StHDR1-cYFP/nYFP, and cYFP/nYFP were regarded as negative controls, respectively. Images were photographed under a fluorescent inverted microscope. Scale bars = 20 μm. (**C-E**) LCI assays were performed to further prove the interactions between StHDR1-Nluc and StDA1-Cluc. Co-expression of StDA1-Cluc/Nluc, StHDR1-Nluc/Cluc, and Nluc/Cluc were regarded as negative controls, respectively. The activity of LUC was measured by chemiluminescence instrument, and the luminescence was measured by plant living imaging system. (**F**) Pull down of StDA1 with StHDR1. GST-StDA1 fusion protein or GST control interaction with His-StHDR1 using anti-His immunoblotting.

Although StHDR1 is a vital enzyme in the aurantio-obtusin biosynthesis pathway in *S.tora* seeds, the genetic regulatory network of StHDR1-mediated aurantio-obtusin biosynthesis remains to be elucidated. To further characterize the effect of StHDR1 in the accumulation of aurantio-obtusin in *S.tora* seeds, overexpression, antisense overexpression of StHDR1 and EV lines were obtained (Figure S5). At 4 months growth stage, we found that the plant height and width of StHDR1OE lines significantly increased while a significant decrease in StDA1AE lines height and width was observed compared to those of EV lines (Figure 5A and Figure S6). At the pod-bulging stage of *S.tora*, the length of pods and seeds were slightly higher in StHDR1OE than in EV lines, on the contrary, they were slightly lower in StDA1AE lines than in EV lines (Figure 5B, C and Figure S6). However, seeds width did not show significant differences in StHDR1OE, StHDR1AE and EV lines (Figure5C and Figure S6). In conclusion, StHDR1 positively regulates the growth and seed formation of *S.tora* plants.

**Fig. 5.**
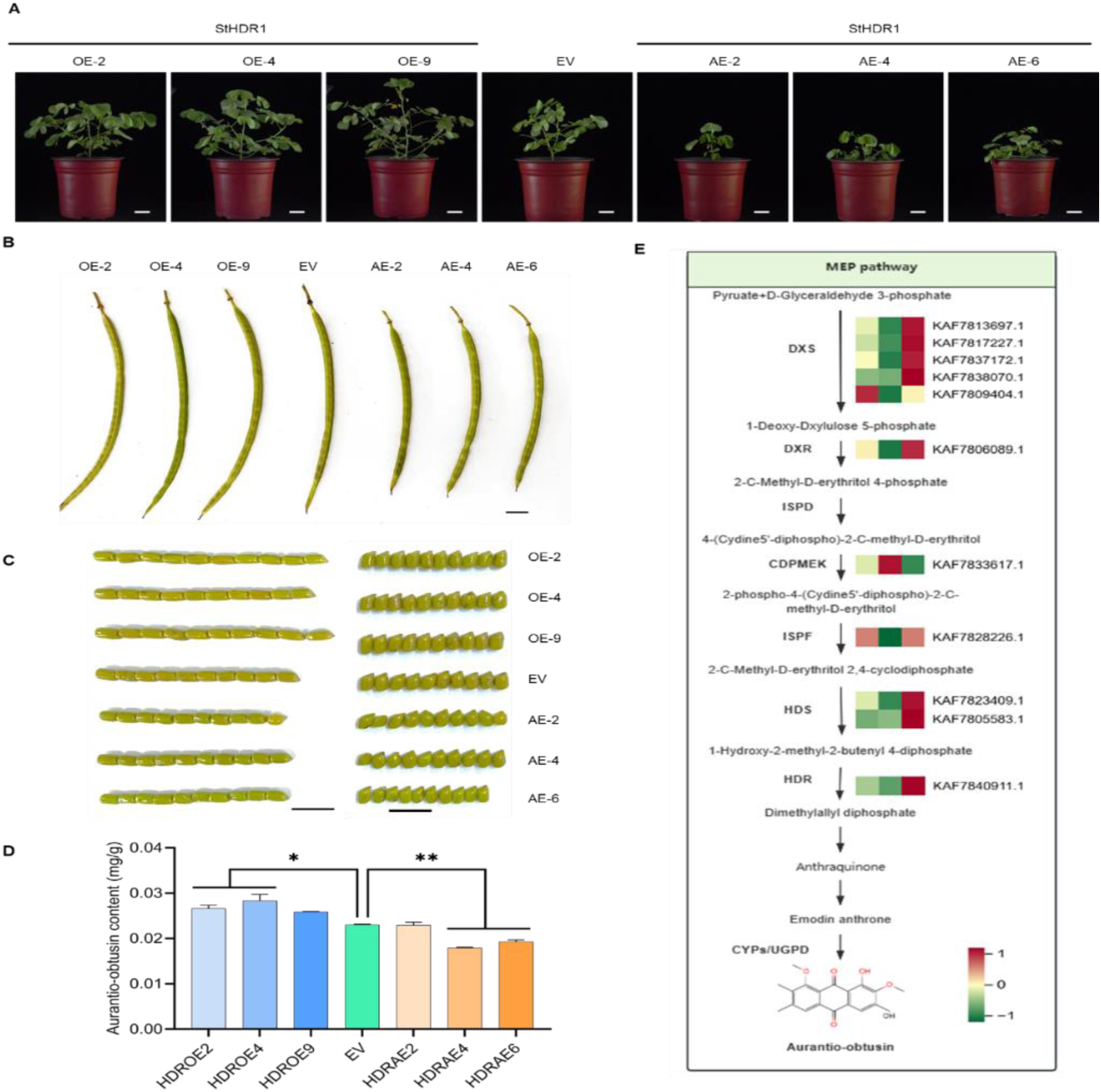
Expression of StHDR1 in different tissues and determination of plant growth and orange determinant content in transgenic lines of StHDR1. (**A**) The plant phenotypic analysis of overexpression and antisense overexpression of StHDR1 lines, scale bars = 5 cm. (**B and C**) The analysis of pod and seed size of overexpression and antisense overexpression of StHDR1 lines, scale bars = 1 cm. (**D**) The aurantio-obtusin content of overexpression, antisense overexpression of StDA1, and EV plant lines. Bars are means ± SD. Statistical analysis was carried out using one-way ANOVA with multiple comparisons, and significant differences were uncovered compared to EV (*, P < 0.05; **, P < 0.01). (**E**) The expression levels of related genes involved in aurantio-obtusin biosynthesis in StHDR1 transgenic plants. The heat map from left to right is empty vector, StHDR1AE, and StHDR1OE lines. The expression level of the gene in the empty vector plant was used as a reference. Red indicates that the gene expression is high, and green indicates that the gene expression is low. DXS, 1-Deoxy-D-xylulose-5-phosphate synthase; DXR, 1-Deoxy-D-xylulose-5-phosphate reductoisomerase; ISPD, 2-C-Methyl-D-erythritol 4-phosphate cytidylyltransferase; CDPMEK, 4-Diphosphocytidyl-2-C-methyl-D-erythritol kinase; ISPF, 2-C-Methyl-D-erythritol 2,4-cyclodiphosphate Synthase; HDS, (E)-4-Hydroxy-3-methylbut-2-enyl-diphosphate synthase HDR, 4-Hydroxy-3-methylbut-2-enyl diphosphate reductase; IPPS, Isopentenyl-diphosphate delta-isomerase.

Furthermore, we found that StHDR1 mainly drives its function in the cell membrane, cytoplasm and nucleus (Figure S7). And its expression is higher at the late stage of *S.tora* seed maturation (Figure S8), which was opposite to StDA1. Subsequently, the content of aurantio-obtusin in StHDR1OE, EV and StHDR1AE lines was analyzed by HPLC. The results showed that aurantio-obtusin content was remarkably higher in StHDR1OE lines and significantly lower in the SHDR1AE lines compared with EV lines (Figure 5D). Meanwhile, we used RT-qPCR to detect the changes of the changes of synthase genes in the MEP pathway at the transcriptional level in StHDR1 transgenic plants. The results revealed that the expression trends of most genes were consistent with those of StHDR1, suggesting that StHDR1 positively regulated the accumulation of aurantio-obtusin in *S.tora* seeds (Figure 5E). The SHDR1 results are contrary to the results of StDA1, suggesting that StHDR1 and StDA1 have opposite functions in the growth regulation, seed formation, and aurantio-obtusin biosynthesis in *S.tora*.

### StDA1 can modulate the stability of StHDR1

It is well known that the UIM structural domain of AtDA1 is mainly involved in combining with ubiquitin(Li et al, 2008). Therefore, we examined the ability of StDA1 to bind ubiquitin. We ligated the full-length sequence of StDA1 and a truncated sequence without two UIMs (StDA1^delUIMs^) to glutathione transferase (GST) fusion proteins, and ubiquitin was expressed as an HA fusion protein. While the GST and the StDA1^delUIMs^-GST did not bind to HA-ubiquitin, StDA1-GST was able to bind to HA-ubiquitin, suggesting that the UIMs of StDA1 are required for binding ubiquitin (Figure 6A). In addition, we investigated whether StDA1 can undergo ubiquitination modification in plants. Total proteins from plants containing StDA1 and GFP were extracted and immunoprecipitated in the presence of MG132 proteasome inhibitor. The results showed that a significant level of ubiquitination occurred in StDA1 plants compared with GFP plants ((Figure 6B)). The above findings implied that the StDA1 protein may be a ubiquitin receptor.

**Fig. 6.**
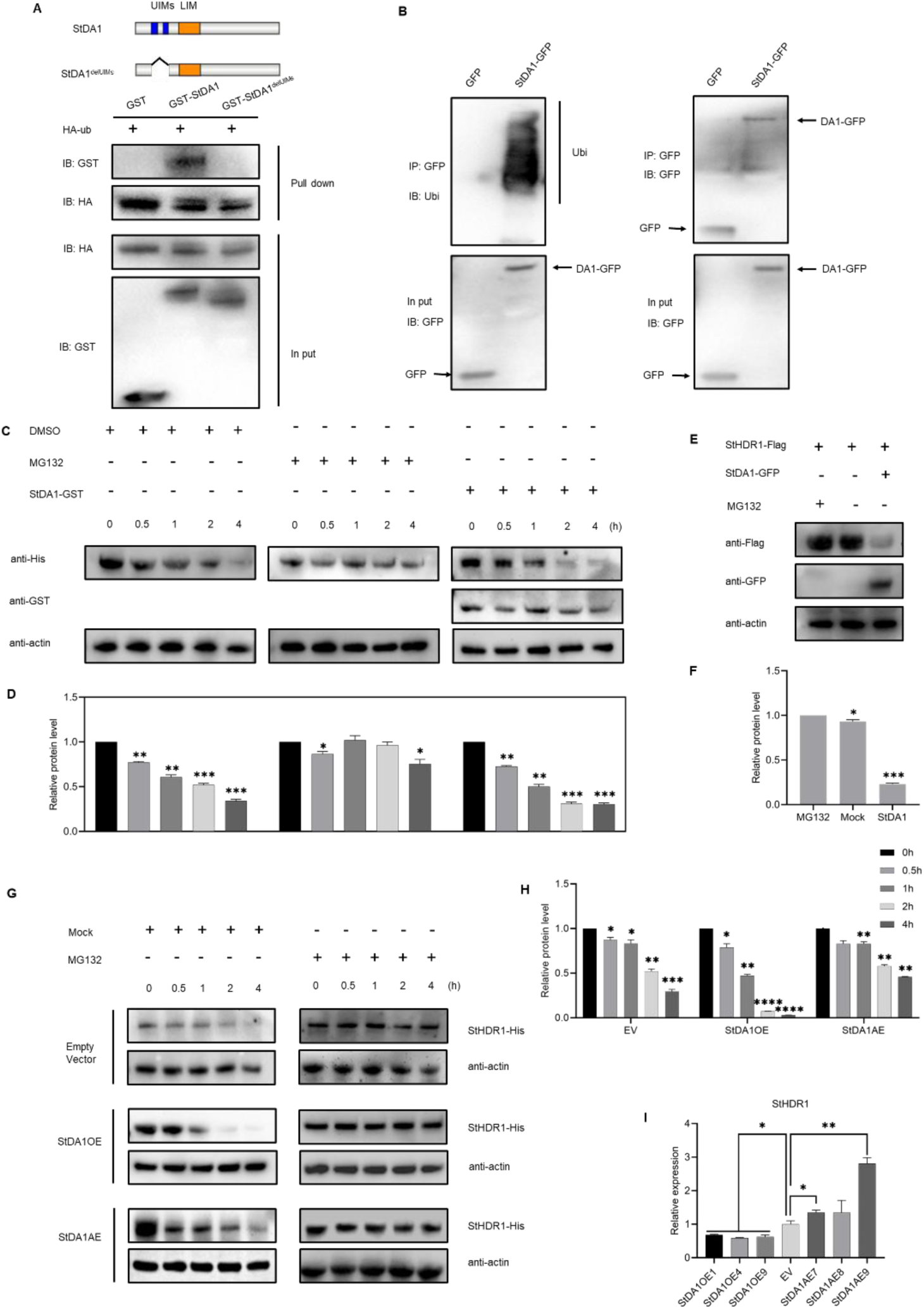
StDA1 modulates StHDR1 stability. (**A) UIMs of StDA1 interact with ubiquitin in vitro.** StDA1-GST, StDA1**^delUIMs^**-GST fusion protein or GST control interaction with HA-StHDR1 using anti-GST immunoblotting. **(B)** Ubiquitination analysis in vivo. Immunoprecipitated proteins were measured by immunoblotting (IB) using anti-ubiquitin and anti-GFP antibodies (**C and D**) StDA1 diminishes the protein stability of StHDR1. Equal amounts of recombinant StDA1-GST and StHDR1-His proteins were incubated with total protein extracts from leaves of 3-month-old *S.tora* for 0, 0.5, 1, 2 and 4 hours without or with 50 mM MG132 added. Total proteins were analyzed by IB using anti-His antibody to quantify the accumulation of StHDR1. Loading control using actin. (**E and F**) StHDR1 protein stability is decreased by StDA1 in vivo. An equal amount of StHDR1-FLAG and StDA1-GFP recombinants were co-transfected into tobacco. StHDR1-Flag and StDA1-GFP protein abundance as determined using anti-Flag and anti-GFP antibodies by IB. Loading control using actin. (**G and H**) StHDR1 protein abundance in empty vector (EV) and StDA1OE/AE transgenic plants. Total protein extracts from leaves of 3-month-old EV and StDA1OE/AE transgenic plants. Recombinant StHDR1-His proteins were incubated in total protein (different genotypes) with or without MG132 for the indicated intervals. StHDR1 accumulation was determined by IB. (**I**) RT–qPCR analysis of the StHDR1 gene expression patterns in EV and StDA1OE/AE transgenic plants. Quantification of protein levels using ImageJ software. Bars are means ± SD. Statistical analysis was carried out using one-way ANOVA with multiple comparisons, and significant differences were uncovered compared to control (*, P < 0.05; **, P < 0.01, ***, P < 0.001, ****, P < 0.0001).

The ubiquitin receptor DA1 is thought to take part in the protein degradation of ubiquitin-mediated interacting proteins(Li et al, 2008; Xia et al, 2013). The ubiquitin proteasome can often be responsible for determining the degradation of numerous proteins. Therefore, to understand how StDA1 affects the function of StHDR1, when the StHDR1-His recombinant protein was incubated with total protein extracted from EV plants, rapid degradation of the StHDR1 protein was observed. However, when it was co-incubated with the recombinant protein StDA1-GST promoted the degradation of StHDR1, which was repressed by the 26S proteasome inhibitor MG132 (Figure 6C and D). It was shown that StDA1 is essential for the stability of the StHDR1 protein.

Meanwhile, we tested the stability of the StHDR1 protein in a tobacco transient expression system. After the StHDR1-Flag construct was transfected into tobacco leaves alone, it was found that StHDR1 accumulated to higher levels after MG132 treatment compared with no treatment. However, when the StHDR1-Flag construct was mixed with StDA1-GFP and transferred into tobacco leaves, StHDR1 abundance became lower (Figure 6E and F). This is in accordance with the outcomes of in vitro cellular degradation experiments.

Moreover, to further assess the stability of the StHDR1 protein, we measured StHDR1 protein levels in StDA1OE, StDA1AE and EV lines. Recombinant StHDR1-His protein was incubated in cell lysates of EV, StDA1OE, and StDA1AE transgenic plants with or without the addition of MG132, respectively, and then was quantified by immunoblotting. Compared with EV plants, StHDR1-His protein showed greater stability in protein extracts from StDA1AE transgenic plants and was more easily degraded in protein extracts from StDA1AE transgenic plants (Figure 6G and H), suggesting that StDA1 is able to contribute to the ubiquitin proteasome-mediated degradation of the StHDR1 protein in *S.tora*. We further examined the abundance of StHDR1 transcripts in EV, StDA1OE, and StDA1AE transgenic plants. The abundance of StHDR1 transcripts accumulated at a higher level in StDA1AE transgenic plants and at a lower level in StDA1OE plants compared with EV plants (Figure 6I). Therefore, we considered that StDA1 influences the stability of StHDR1 in *S.tora*.

### StTCP4.1 can facilitate the interaction between StDA1 and StHDR1

To further investigate how the StDA1 influences the stability of StHDR1, we found that the transcription factor StTCP4.1 can interact with StDA1 from the StDA1 pull down-MS results. Alternatively, we identified the ability of StTCP4.1 to negatively regulate the growth of *S.tora*, seed formation, and aurantio-obtusin accumulation (Figure S9-S11) by observing the phenotypes of StTCP4.1 overexpression, antisense overexpression, and EV lines and performing HPLC. To further verify the interaction between StDA1 and StTCP4.1, we performed Y2H, BiFC and pull down assays.Y2H assays showed that yeast cells co-expressing StDA1-BD and StTCP4.1-AD could grow on SD/-Leu-Trp-His-Ade medium containg 200 ng/ml AbA (Figure S12A), indicating that StTCP4.1 could interact with StDA1. BiFC analysis displayed that obvious YFP fluorescence could be observed on tobacco leaves with StDA1-nYFP / StTCP4.1-cYFP co-expression (Figure S12B). Consistent with the BiFC assay, LCI assays were conducted to further confirm higher LUC activity and stronger luminescence signal in the StDA1-Cluc/StTCP4.1-Nluc co-expressed tobacco leaves but not in the negative control (Figure S12C-E), suggesting that StDA1 can interact with StTCP4.1in vivo. Then, we performed pull down experiments with purified recombinant GST-tagged StDA1 (StDA1-GST) and His-tagged StTCP4.1 (StTCP4.1-His) and found that the StTCP4.1-His protein could pull down StDA1-GST, which indicated that StDA1 physically interacts with StTCP4.1 in vitro (Figure S12F).

Since StTCP4.1 is a member of the TCP transcription factors, we speculated whether StTCP4.1 could regulate StDA1 at the transcriptional level in addition to the protein level. We found that there is no binding site of the StTCP4.1 transcription factor on the promoter of StDA1 by analyzing the cis-elements of the 1500 bp promoter sequence of StDA1 (Table S3). Meanwhile, we further verified using Y1H assay, which illustrated that yeast cells co-transfected with the promoter sequence of StDA1 and StTCP4.1-AD also failed to grow on SD/-Ura containing Aureobasidin A (AbA) medium, indicating that StTCP4.1 could not interact with the promoter of StDA1 (Figure S13). In summary, the transcription factor StTCP4.1 can only interact with StDA1 at the protein level.

We analyzed the interaction regions of StDA1 with StHDR1 and StTCP4.1 by Y2H assays. The results showed that StDA1-LIM containing intermediate region LIM interacted with StHDR1, while StDA1-UIMs containing N-terminal UIMs interacted with StTCP4.1, and StDA1-C containing C-terminal region was not able to interact with the above two proteins (Figure 7A), indicating that the domains of UIMs and LIM of StDA1 protein are important segments for its interaction with other proteins. Meanwhile, since StHDR1 and StTCP4.1 have different regions of interactions with StDA1, StTCP4.1 and StDA1-StHDR1 modules may form a complex driving function.

**Fig. 7.**
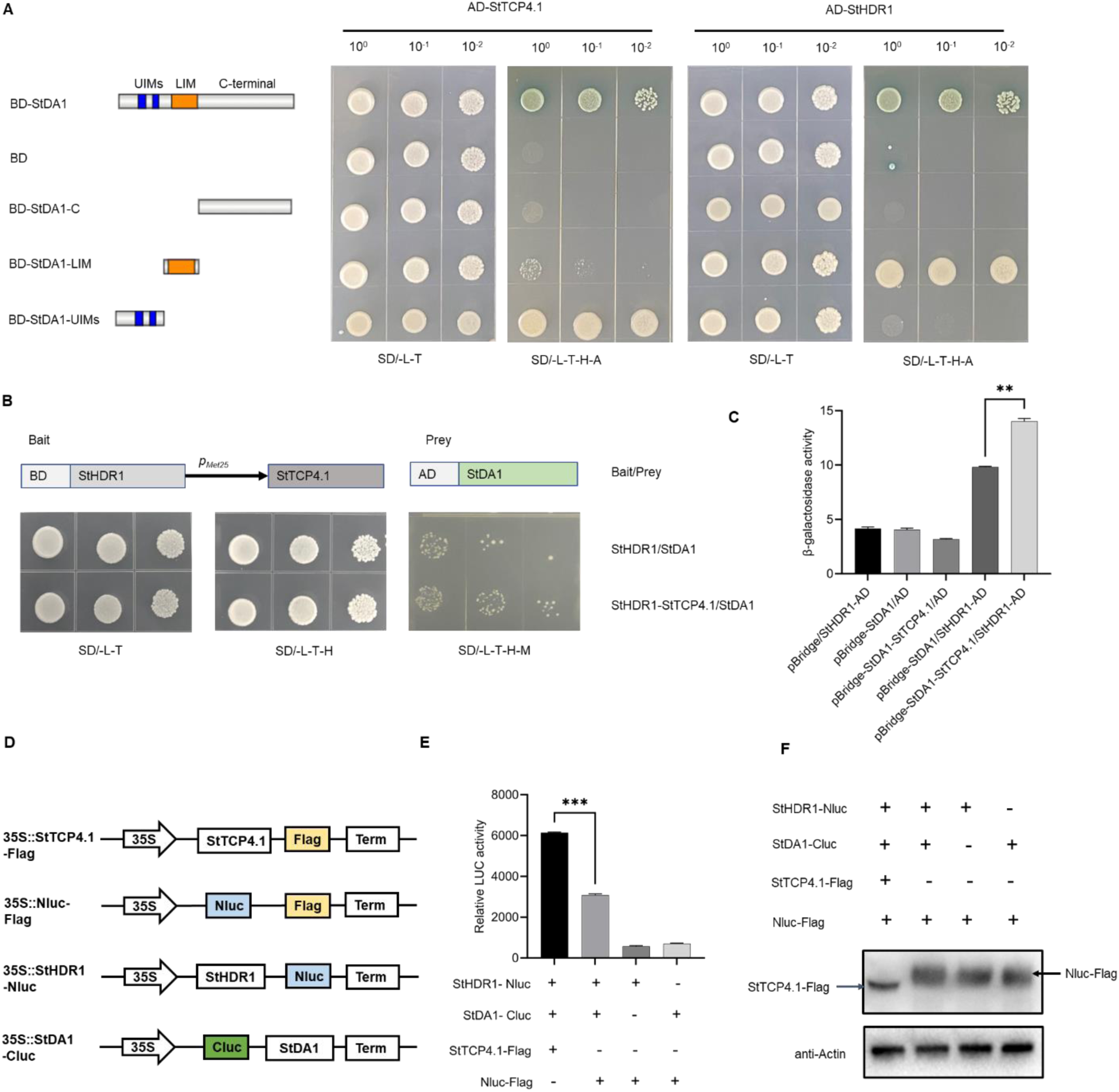
StTCP4.1 influences StDA1 interactions with StHDR1. (**A**) Interaction of different regions of StDA1 with StHDR1/StTCP4.1 in Y2H assays. The StDA1 sequence is divided into three parts with two UIM motifs at the N-terminus, a LIM structural domain in the middle region, and a C-terminal region. The yeast cells were observed after 4 days of growth on SD/-Trp-Leu and SD/-Trp-Leu-His-Ade medium containing AbA and X-a-gal. (**B**) StTCP4.1 affects the interaction between StDA1 and StHDR1 in Y3H. PMet25 is an inducible promoter that initiates expression of the intermediate protein StTCP4.1. The yeast cells were observed after 4 days of growth on SD/-Trp-Leu, SD/-Trp-Leu-His and SD/-Trp-Leu-His-Met medium containing 3-AT. (**C**) Analysis of β-galactosidase activity of yeast in (B). Yeast colonies produce β-galactosidase, which is capable of breaking down o-nitrophenyl-β-D-galactopyranoside (ONPG) to produce the yellow o-nitrophenol (ONP). (**D and E**) StTCP4.1 affects the interaction between StDA1 and StHDR1 in LCI assays. The activity of LUC was measured by chemiluminescence instrument. Bars are means ± SD. Statistical analysis was carried out using Student’s t-test, and significant differences were uncovered (**, P < 0.01, ***, P < 0.001). (**F**) Analysis of protein expression of tobacco in LCI assay. StTCP4.1-Flag and Nluc-Flag proteins were detected by anti-Flag antibody.

To understand how StTCP4.1 forms a complex with StDA1 and StHDR1, we used Y3H assays to investigate whether StTCP4.1 affects the protein interactions of StDA1 and StHDR1. When all three proteins (StTCP4.1, StDA1 and StHDR1) were expressed simultaneously, it was clearly found that yeast cells containing the three proteins grew stronger on the culture medium than yeast containing only two proteins (StDA1 and StHDR1) (Figure 7B). Subsequently, the β-galactosidase assay in this study also confirmed that the β-galactosidase activity of yeast colonies containing three proteins (StTCP4.1, StDA1 and StHDR1) was stronger than that of yeast colonies containing only two proteins (StDA1 and StHDR1) (Figure 7C). Meanwhile, LCI assays in tobacco showed that the LUC activity was higher when StTCP4.1-Flag was co-expressed with StHDR1-Nluc and StDA1-Cluc than when StHDR1-Nluc and St-DA1-Cluc were co-expressed, suggesting that StTCP4.1 promotes the interaction of StDA1 and StHDR1 (Figure 7D-F).

Furthermore, we analyzed the expression of StTCP4.1 in StDA1 and StHDR1 transgenic plants, and we found that StTCP4.1 expression was up-regulated in StDA1OE lines and down-regulated in StDA1AE lines (Figure S14). However, the expression of StTCP4.1 in StHDR1 transgenic plants was completely in contrary to that in StDA1 transgenic plants, suggesting that StTCP4.1 has the same expression pattern with StDA1 and the opposite expression pattern with StHDR1.

### MeJA treatment can inhibit the promoting effect of StTCP4.1 on StDA1-StHDR1 module

It is well known that jasmonic acid and its derivative methyl jasmonate (MeJA) usually have a promoting effect on the accumulation of secondary metabolites in plants(Liu et al, 2023). In *S.tora* hairy roots, jasmonic acid has been proven to promote the accumulation of anthraquinones in early studies(Yang et al, 2005). Meanwhile, we found that MeJA was able to affect the accumulation of aurantio-obtusin in *S.tora* seeds (Figure S15), however, it is not clear how MeJA and the StDA1-StHDR1 module together affect the regulatory network mechanism of aurantio-obtusin biosynthesis in *S.tora* seeds. It has been demonstrated in our previous studies that StTCP4.1 is involved in the MeJA-regulated pathway in *S.tora* and its expression is repressed by MeJA(Liu et al, 2022). Therefore, we speculated whether the StDA1-StHDR1 would also response to MeJA treatment. Therefore, *S.tora* leaves were treated with 250 uM MeJA, and it was found that after 4 h, the expression of StDA1 was significantly down-regulated, whereas StHDR1 expression was significantly up-regulated, indicating that the expression of both StDA1 and StHDR1 responded to MeJA (Figure 8A).

**Fig. 8.**
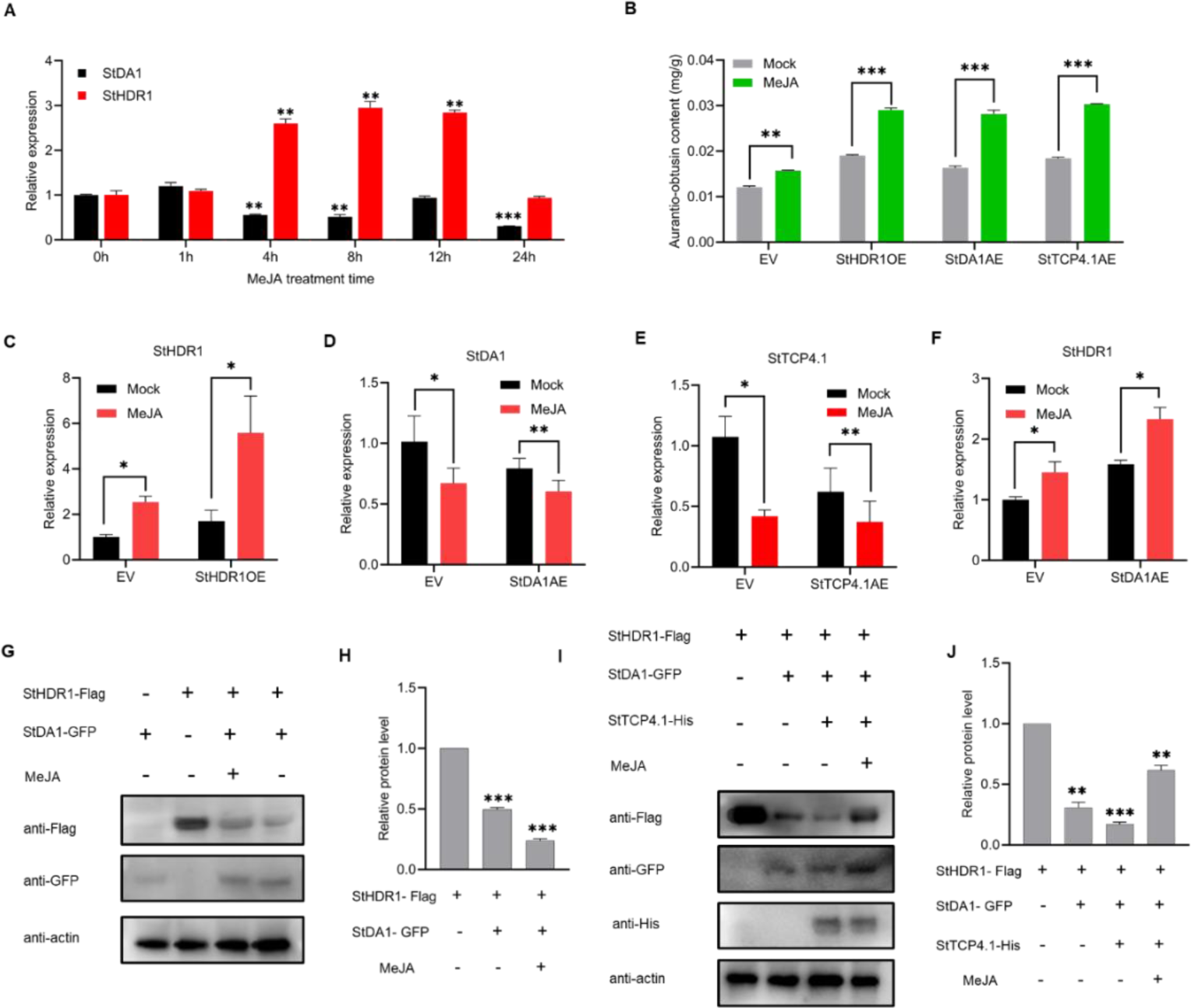
The StDA1-StHDR1 module-driven accumulation of aurantio-obtusin is responsive to MeJA in *S.tora* seeds. (**A**) RT-qPCR analysis of StDA1 and StHDR1 expression under different MeJA treatment time. Bars are means ± SD. Statistical analysis was carried out using the Student’s t-test, and significant differences were uncovered compared to 0 h (**, P < 0.01, ***, P < 0.001). (**B**) The aurantio-obtusin content in MeJA-treated and untreated control (EV), StHDR1OE, and StDA1AE lines using HPLC. (**C-E**) RT-qPCR analysis of StHDR1/StDA1/StTCP4.1 expression in MeJA-treated and untreated control (EV) and StHDR1OE/StDA1AE/StTCP4.1AE lines. (**F**) RT-qPCR analysis of StHDR1 expression in MeJA-treated and untreated control (EV) and StDA1AE lines. (**B-F**) Bars are means ± SD. Statistical analysis was carried out using the Student’s t-test, and significant differences were uncovered compared to mock (*, P < 0.05, **, P < 0.01, ***, P < 0.001). (**G and H**) MeJA treatment alleviated the degradation of StHDR1 by StDA1. An equal amount of StHDR1-Flag and StDA1-GFP recombinants were co-transfected into tobacco. After 48 hours of 16 light/8 dark treatment, the leaves were treated with 100 uM MeJA for 4 hours. StHDR1-Flag and StDA1-GFP protein abundance as determined using anti-Flag and anti-GFP antibodies by IB. (**I and J**) MeJA inhibited the promoting effect of StTCP4.1 on StDA1-StHDR1 module. An equal amount of StHDR1-Flag, StTCP4.1 His and StDA1-GFP recombinants were co-transfected into tobacco. After 48 hours of 16 light/8 dark treatment, the leaves were treated with 100 uM MeJA for 4 hours. StHDR1-Flag, StTCP4.1-His and StDA1-GFP protein abundance as determined using anti-Flag, anti-His and anti-GFP antibodies by IB. Loading control using actin. Quantification of protein levels using ImageJ software. Bars are means ± SD. Statistical analysis was carried out using one-way ANOVA with multiple comparisons, and significant differences were uncovered compared to EV (*, P < 0.05; **, P < 0.01, ***, P < 0.001).

Subsequently, fresh seeds of EV, StDA1AE, StTCP4.1AE, and StHDR1OE transgenic plants were treated with 250 uM MeJA, and HPLC results revealed that MeJA promoted aurantio-obtusin accumulation in EV, StDA1AE, StTCP4.1AE, and StHDR1OE transgenic plants (Figure 8B). We further examined the effect of MeJA treatment on the abundance of StDA1, StTCP4.1 or StHDR1 transcripts in StDA1AE, StTCP4.1AE or StHDR1OE transgenic plants. MeJA promoted StHDR1 expression in StHDR1OE plants, and repressed StDA1/StTCP4.1 expression in StDA1AE/StTCP4.1AE plants compared with untreated plants (Figure 8C-E).

Additionally, we tested the abundance of StHDR1 transcripts in MeJA-treated EV and StDA1-OE transgenic plants. The abundance of StHDR1 transcripts accumulated at significantly higher levels in StDA1OE transgenic plants compared with the control (Figure 8F). We further determined whether MeJA affected the stabilization of StHDR1 protein by StDA1 through a tobacco transient expression system. The results revealed that the abundance of StHDR1 was low in the extracted cell lysates after the StHDR1-Flag construct was transferred into tobacco leaves mixed with StHDR1-GFP. In contrast, StHDR1 accumulated to slightly higher levels after MeJA treatment (Figure 8G and H). On this basis, we simultaneously transfected StTCP4.1-His, StHDR1-Flag and StHDR1-GFP into tobacco and found that StTCP4.1 accelerated the degradation of StHDR1 by StDA1, however, the degradation level of StHDR1 was alleviated after Me JA treatment (Figure 8I and J). These results suggest that MeJA inhibited the promoting effect of StTCP4.1 on StDA1-StHDR1 module.

## Discussion

### StDA1 negatively regulates the accumulation of aurantio-obtusin through seed formation

*S.tora* is the most common herb in the legume family and is used as medicine especially its seeds. Modern pharmacological studies have found that *S.tora* has plenty of therapeutic effects, such as antifertility, antihypertensive, anti-inflammatory, antifungal, anti-glycation, anti-hyperglycemic, antibacterial activity (Ahmad et al, 2018). The main active ingredients of *S.tora* are anthraquinones, such as aurantio-obtusin, chrysophanol, emodin, rhein and obtusifolin (Guo et al, 2017). However, aurantio-obtusin is a unique component in *S.tora* seeds (Figure S2). With the wide application of *S.tora* seeds in the production of Chinese patent medicine, clinical formulas, health care products and food industry, the demand for *S.tora* seed is gradually increasing every year. In recent years, most studies on *S.tora* have focused on the key enzymes within aurantio-obtusin biosynthesis pathway, whereas the understanding of how seed formation and aurantio-obtusin accumulation are regulated in terms of post-translational modifications of the protein is still poor. In the present study, we detected a ubiquitin receptor, StDA1 whose expression reaches its lowest level at the late stage of seed maturation; then we showed that this receptor has a negative regulatory role in seed formation and aurantio-obtusin accumulation through the ubiquitin-proteasome pathway in *S.tora* (Figure 1B and C).

The ubiquitin receptor DA1 is one of the components of the ubiquitin-proteasome pathway, and it has been previously shown that DA1 can negatively regulate seed size by restricting the proliferation of matricellular cells in *A.thaliana*, rapeseed and wheat(Li et al, 2008; Liu et al, 2020; Wang et al, 2017). Besides, overexpression of mutated AtDA1 in rapeseed also leads to the enlargement of organs such as flowers, cotyledons, seeds, leaves, and leaf sheaths(Wang et al, 2017). And in maize, it negatively regulates seed size by inhibiting sugar transport into the maize endosperm(Xie et al, 2018). These findings point to the fact that DA1 is important in the molecular mechanism regulating seed size. In our results, overexpression of StDA1 leads to a decrease in pods number, seeds size, and aurantio-obtusin accumulation whereas StDA1 antisense overexpression increased the number of pods, the size of seeds, and the content of aurantio-obtusin (Figure 2 and 3), indicating that StDA1 regulates secondary metabolites accumulation in plants except for the regulation of plant growth and seed formation, which provides a solid theoretical basis for the regulatory mechanism of DA1 in the accumulation of secondary metabolites in plants.

### StDA1 regulates the stability of StHDR1 protein

Although there are relatively few studies on the effect of DA1 in plant secondary metabolites biosynthetic pathways. However, in poplar, WOX4 is the target of DA1 ubiquitination. The ubiquitin-proteasome pathway can regulate the stability of the WOX4 protein to regulate cambium cell division in the formative layer and xylem formation(Tang et al, 2022). In *A.thaliana*, many key enzymes have also been detected as targets of ubiquitination in the phenylpropanoid pathway that regulates anthocyanin biosynthesis, including PAL, C4H, HCT, CAD, and OMT1. In the present study, HDR1 is the terminal enzyme of the MEP pathway that regulates the biosynthesis of aurantio-obtusin in *S.tora* and was found to be a novel target of DA1 ubiquitination. StDA1 regulates the degradation of the StHDR1 protein, which provides a new idea for the regulation of HDR1 at the protein level.

In this study, the expression level of StDA1 was relatively low at the late stage of seed maturation, while StHDR1 was the opposite (Figure 1B and Figure S8). These two proteins can interact with each other (Figure 4), suggesting that they may be linked during seed formation. The presence of two UIM motifs on the StDA1 sequence that are essential for ubiquitin binding (Figure 6A), as well as the fact that StDA1 can be ubiquitinated in plants (Figure 6B), suggests that StDA1 may be a ubiquitin receptor involved in the ubiquitin-mediated protein degradation of interacting proteins. In vitro cell-free protein degradation assays showed that the degradation rate of StHDR1 protein in StDA1OE plants was faster than that in StDA1AE plants (Figure 6C-H). We concluded that StHDR1 and StDA1 have antagonistic effects during seed formation, and StDA1 mediates seed formation and the accumulation of aurantio-obtusin in seeds by regulating the stability of StHDR1. Consistent with this hypothesis, the relative abundance of StHDR1 transcripts was up-regulated in StDA1AE plants and down-regulated in StDA1OE transgenic plants (Figure 6I), indicating the role of StDA1 in StHDR1 degradation.

### The StTCP4.1-StDA1-StHDR1 complex is involved in aurantio-obtusin biosynthesis

Based on the results of our previous experiments, StDA1 and StHDR1 have opposite roles in regulating the biosynthesis of aurantio-obtusin (Figure 2L and Figure 4D). Interestingly, this study found that StDA1 and StHDR1 also have opposite roles in response to JA (Figure 8A). Similarly, we found that after MeJA treatment of the seeds of StDA1AE and StHDR1OE transgenic plants, the expression of StHDR1 in StHDR1OE plants was promoted, and the expression of StDA1 in StDA1AE plants was down-regulated. The content of aurantio-obtusin in both transgenic plants was significantly elevated (Figure 8C and D), further indicating that MeJA mediates aurantio-obtusin biosynthesis regulation mechanism by StDA1 and StHDR1. We also found that StDA1 and StHDR1 interacted at the protein level (Figure 4) and could degrade StHDR1 protein (Figure 6) without MeJA treatment. The degradation of StHDR1 protein by StDA1 was inhibited with MeJA treatment (Figure 8G and H). These results demonstrate that MeJA mediates the regulation mechanism of aurantio-obtusin biosynthesis by StDA1 and StHDR1 through inhibiting StHDR1 protein degradation by StDA1.

In plants, TCP transcription factors can directly or indirectly interact with key genes in the JA signaling pathway and participate in secondary metabolism accumulation. In *Artemisia annua*, the JA signaling repressor AaJAZ8 interacts with the AaTCP14-AaORA complex to regulate the activity of the DBR2 promoter, which affects artemisinin biosynthesis(An et al, 2018). In *S.tora*, StTCP4.1 can play a role in the JA signaling pathway by interacting with StMYC2a(Liu et al, 2022). Subsequently, we identified another link between the StDA1-StHDR1 complex and the JA signaling pathway, that is, JA-responsive StTCP4.1. In this study, StDA1 interacts with StTCP4.1 and StHDR1, respectively, and StTCP4.1 can bind to the UIMs structural domains of StDA1, and StHDR1 can bind to the LIM structural domains of StDA1, resulting in the formation of a StTCP4.1-StDA1-StHDR1 complex (Figure 7A). It was also found that StTCP4.1 could promote the interaction of StDA1 with StHDR1 (Figure 7B-F), as well as being able to promote the degradation of StHDR1 by StDA1 (Figure 8I-J). However, the degradation of StHDR1 protein was inhibited after MeJA treatment (Figure 8I-J).

Based on the above experimental results, we proposed a hypothetical model to elucidate the mechanism of aurantio-obtusin regulation in *S.tora* by the MeJA-mediated StDA1-StHDR1 module (Figure 9). In the absence of jasmonic acid, free StTCP4.1 in the cells formed a complex with StDA1 and StHDR1, which facilitated the interaction of StDA1 with the key enzyme StHDR1, as well as the degradation of StHDR1, ultimately leading to a decrease in aurantio-obtusin accumulation. In the presence of jasmonic acid, jasmonic acid promoted the interaction of StMYC2 and StTCP4.1, reduced free StTCP4.1 in the cell, which in turn reduced the promotion of StDA1-StHDR1 interaction by StTCP4.1, attenuated the degradation of StHDR1, released more StHDR1 protein, and ultimately promoted aurantio-obtusin accumulation. In summary, our study not only reveals that StDA1 can directly regulate the protein stability of key enzymes in the secondary metabolic pathway, but also elaborates on the regulation of the StDA1-StHDR1 module by MeJA-responsive StTCP4.1, which will provide a solid theoretical foundation for further research on the mechanism by which phytohormones and DA1 synergistically regulate the accumulation of plant secondary metabolites.

**Fig. 9.**
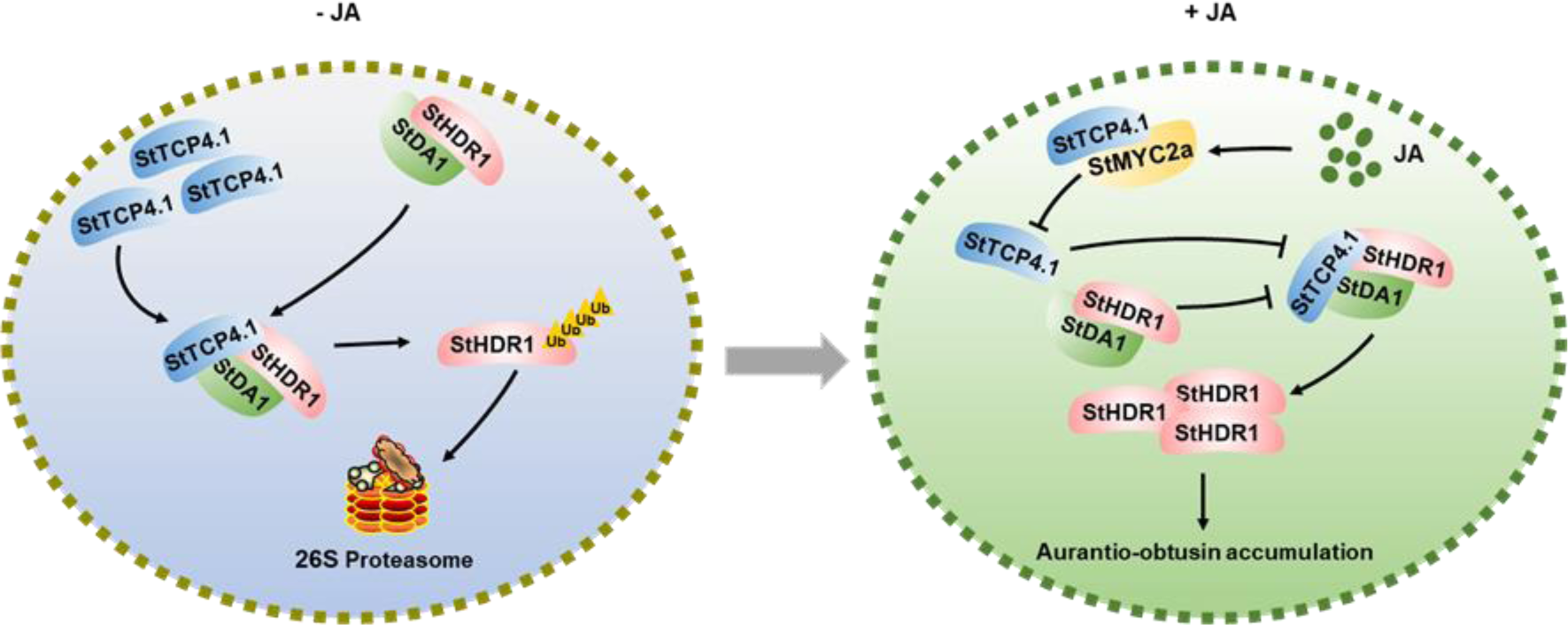
Model of the MeJA-mediated StDA1-StHDR1 module regulating the accumulation of aurantio-obtusin in *S.tora*. In the absence of jasmonic acid, StTCP4.1 formed a complex with StDA1 and StHDR1, which promoted the interaction of StDA1 with the key enzyme StHDR1 and degradation of StHDR1, while decreasing the accumulation of aurantio-obtusin. In the presence of jasmonic acid, jasmonic acid promotes the interaction of StTCP4.1 with StMYC2a, which in turn reduces the facilitating effect of StTCP4.1 on the StDA1-StHDR1 interaction, attenuates the degradation of StHDR1, releases more StHDR1 proteins, and ultimately facilitates the accumulation of aurantio-obtusin. Arrows indicate promotion and bars (T) indicate inhibition.

## Materials and methods

### Plant materials and growth conditions

The seeds of *S.tora* were provided by Shaanxi Tianshili Botanical Pharmaceutical Co. The seeds were soaked in a thermostatic water bath at 50°C for 5-6 h, planted in pots, and grown at 25°C, 16 h light/8 h dark. The leaves of *S.tora* grown for about 1 month were collected for RNA extraction and reverse transcribed into cDNA. The CDS of StDA1, StHDR1, StTCP4.1 were amplified according to the primer sequences (Table S4). And the three genes were ligated to pCAMBIA1300-mCherry by forward and reverse, respectively, in order to obtain overexpression plants and antisense overexpression plants. Freshly sprouted *S.tora* seeds were placed in a centrifuge tube and *Agrobacterium* resuspension was poured until the seeds were completely submerged. The vacuum pump was used to evacuate the seeds. When the vacuum pressure reached 0.08 MPa, the vacuum was stopped and timed for 5 min, and after the timer ended, the pressure was quickly released and repeated 3 times. The germinated seeds were transplanted to the soil and cultivated in the dark for 24 h, and then restored to 16-h light/8-h dark incubation. Furthermore, tobacco seedlings were grown under 16 h light/8 h dark conditions.

### RT-qPCR analysis

The roots, stems, leaves, and flowers of *S.tora* were collected at the flowering stage, and the seeds were collected during the podding stage, with three replicates of each tissue being collected. The leaves of *S. tora* were sprayed with 250 uM MeJA solution, and samples were collected at 0 h, 1 h, 4 h, 8 h, 12 h, and 24 h after treatment, respectively. For each treatment, three biological replicates were set up and after about one months and the leaves were collected. The Steady Pure plant RNA extraction kit (Accurata Biotech, Hunan, China) was utilized to extract RNA from the samples. The 5X Evo M-MLV RT Master Mix and SYBR® Green Premix Pro Taq HS qPCR Kit (Accurata Biotech, Hunan, China) were used to perform RNA reverse transcription and RT-qPCR, respectively. The specific primers designed by Primer3 are shown in Table S4.

### Determination of aurantio-obtusin content

Seeds of *S.tora* were grounded into powder using a pulverizer. 0.1 g of powder was added to 10 ml of extraction solution (methanol: hydrochloric acid = 10:1) and soaked overnight followed by sonication for 2 h. The mixture was centrifuged at 100,00 g for 10 minutes. The supernatant was filtered through a 0.22 μm membrane and analyzed by HPLC.

### Subcellular localization

The CDS of StDA1 and StHDR1 were cloned into pCAMBIA1300-GFP to obtain StDA1/StHDR1-GFP. It was shown that StTCP4.1 is located in the nucleus of *S.tora*. Therefore, we cloned StTCP4.1 into pCAMBIA1300-mCherry as a control(Liu et al, 2022). Equal amounts of StDA1/StHDR1-GFP and StTCP4.1-mCherry of *Agrobacterium* were mixed and injected into tobacco leaves. The results were observed after 2 ∼ 3 days.

### Screening for interacting proteins

Interacting proteins of StDA1 were screened by liquid chromatography-mass spectrometry/mass spectrometry (LC-MS / MS)(An et al, 2023; Dahro et al, 2022). Briefly, total proteins extracted from *S.tora* seeds (wild-type) were incubated with StDA1-His proteins together, and StDA1-His interacting proteins could be screened by the pull down technique, which was detected by utilizing anti-His antibody (Abmart, Shanghai, China). The proteins interacting with StDA1 could be detected by further characterization of the pull down protein complexes using LC-MS/MS.

### Y2H assays

The fragment encoding specific protein domains and the CDS of StDA1 were cloned into the pGBKT7 vector to obtain BD-DA1, BD-DA1-Uim, BD-DA1-Lim, and BD-DA1-C constructs. StHDR1 and StTCP4.1 were introduced into the pGADT7 vector to obtain AD-HDR1 and AD-TCP4.1 constructs, respectively. The AD-HDR1 and AD-TCP4.1 constructs were co-transformed into yeast strain Y2HGold with these BD constructs, respectively. The transformed yeast cells were coated on SD-Trp/-Leu and SD-Trp/-Leu/-His/-Ade media containing 200ng/ml AbA and cultured continuously at 28℃ for 4 days.

### BiFC and LCI assays

For bimolecular fluorescence complementation (BiFC) assays, StDA1 was cloned into pXY106 to produce StDA1-nYFP fusion proteins. StHDR1 and StTCP4.1 were cloned into pXY104 to prepare StHDR1-cYFP and StTCP4.1-cYFP fusion proteins. Equal amounts of StDA1-nYFP and StHDR1/StTCP4.1-cYFP Agrobacterium were co-injected into tobacco leaves and incubated for 2-3 days. YFP fluorescence in tobacco leaves was imaged using a fluorescence inverted microscope.

For luciferase complementation (LCI) assay, StDA1 was cloned into pCAMBIA1300-Cluc to produce StDA1-Cluc fusion proteins. StHDR1 and StTCP4.1 were cloned into pCAMBIA1300-Nluc to prepare StHDR1-Nluc and StTCP4.1-Nluc fusion proteins. Equal amounts of StDA1-Cluc and StHDR1/StTCP4.1-Nluc *Agrobacterium* were co-injected into tobacco leaves and incubated for 2-3 days. The activity of LUC was measured by chemiluminescence instrument, and the luminescence was measured by plant living imaging system. Additionally, to analyze the effect of StTCP4.1 on the interactions of StHDR1 and StDA1. Flag sequence was constructed into pCAMBIA1300-Nluc vector and StTCP4.1 was ligated into pCAMBIA1300-flag vector. The *Agrobacterium* containing StDA1-Cluc, StHDR1-Nluc, and StTCP4.1-Flag/Nluc-Flag were injected into tobacco. The LCI experiment was performed according to the above experimental steps. Subsequently, the above samples were added to 4× SDS loading buffer and boiled in a boiling water bath for 10 min before being subjected to Western Blots with anti-Flag antibody (Abmart, Shanghai, China).

### Pull down assays

The StDA1 was cloned into the pGEX4T-1 vector to obtain the StDA1-GST construct. StHDR1 and StTCP4.1 were introduced into the pCold-TF vector to obtain StHDR1-His and StTCP4.1-His constructs, respectively. StDA1-GST protein was purified with ProteinIso® GST Resin (TransGen, Beijing, China), and StHDR1/StTCP4.1-His protein was purified with ProteinIso® Ni-NTA Resin (TransGen, Beijing, China). The purified StDA1-GST protein was mixed with StTCP4.1/StHDR-His proteins adsorbed on the NiNTA Resin, and 300 ul of pull-down buffer (40 mM HEPES-KOH at pH 7.5, 0.4 M Sucrose,10 mM KC1, 3 mM MgCl_2_, 1 mM EDTA, 1 mM DTT, 0.2% Triton X-100, 100 uM protease inhibitor cocktail) was incubated overnight for 4 ℃. Subsequently, the retained Ni-NTA Resin were washed 3-5 times using PBS buffer and analyzed with anti-His antibody (Abmart, Shanghai, China) and anti-GST antibody (Abmart, Shanghai, China).

### Y3H and β-Gal activity assays

The StHDR1 was cloned into the pGADT7 vector to generate the AD-HDR1 construct. StDA1 and StTCP4.1 fragments were cloned into the two multiple cloning sites of the pBridge vector to construct the bait vector pBridge-StTCP4.1-StDA1. The bait vector pBridge-StTCP4.1-StDA1 was co-transformed with AD-StHDR1 into strain AH109, and the transformed yeast cells were coated with SD-Trp, SD-Trp/-Leu/-His and SD-Trp/-Leu/-His/-Met medium, and cultured continuously at 28 ℃ for 4 days.

β-galactosidase (β**-**Gal) assay is capable of being used to quantify protein interaction. In Y3H assays, the interacting yeast colonies can produce β-galactosidase, which was able to break down the substrate o-nitrophenol-β-galactoside (ONPG) to produce yellow o-nitrophenol (ONP).

### Ubiquitin binding and ubiquitinylation assays

To test the possibility of DA1 binding to ubiquitin, we cloned the deleted UIM sequences (StDA1delUIMs) into pGEX4T-1 to obtain StDA1delUIMs-GST. StDA1delUIMs-GST protein was purified with ProteinIso® GST Resin (TransGen, Beijing, China). The recombinant HA-ubiquitin was mixed with purified GST/ StDA1-GST/StDA1delUIMs-GST proteins adsorbed on the GST Resin, the pull down experiment was performed according to the above experimental steps, and analyzed with anti-GST antibody (Abmart, Shanghai, China) anti-HA antibody (Abmart, Shanghai, China).

The CDS of StDA1 were amplified according to the primer sequences (Table S4), and were ligated to pCAMBIA1300-GFP. *Agrobacterium* containing GFP and StDA1-GFP was transfected into *S.tora* according to the above transgenic technique to obtain *S.tora* plants containing GFP and StDA1-GFP. Total proteins were extracted from *S.tora* leaves of *S.tora* plants containing GFP and StDA1-GFP using buffer (10 mM Tris/HCl (pH 7.5), 1-mM EDTA, 150 mM NaCl, 0.5% Nonidet P-40, 10% glycerol, 100 uM protease inhibitor cocktail, and 50 mM MG132). were carried out according to the instruction manual of GFP-Nanoab-Magnetic (Lablead, Beijing, China), anti-GFP (Abmart, Shanghai, China) and anti-Ubi antibodies (Abmart, Shanghai, China) were used for ubiquitination analysis.

### Cell free protein degradation assay

The cell free protein degradation assay was modified based on previous experiments(Gui et al, 2019). In short, total proteins were extracted from 4-month-old *S.tora* leaves of different genotypes using degradation buffer (25 mM Tris/HCl, pH 7.5, 10 mm NaCl, 10 mM MgCl_2_, 100 uM protease inhibitor cocktail, 5 mM DTT, and 10 mM ATP). The protein concentration of the supernatant was detected using the BCA method. Total protein from *S.tora*, recombinant StHDR1-His protein and 50 mM MG132 were incubated together (at room temperature), and samples were collected at 0, 0.5, 1, 2, and 4 h. The abundance of StHDR1-His was ultimately examined by immunoblotting with an anti-His antibody (Abmart, Shanghai, China).

### Immunoblotting

The CDS of StHDR1 was cloned into pCAMBIA1300-Flag, the CDS of StDA1 was cloned into pCAMBIA1300-GFP, and the CDS of StTCP4.1 was cloned into pCAMBIA1300-HIS to produce StHDR1-Flag, StDA1-GFP, and StTCP4.1-His. To determine whether StTCP4.1 will affect the degradation of StHDR1 protein by StDA1, equal amounts of StHDR1-Flag, StDA1-GFP and StTCP4.1-His *Agrobacterium* were co-injected on tobacco leaves. In addition, to assess the effect of MeJA on the interactions of St TCP4.1, StDA1 and StHDR1, 100 uM MeJA was used to spray evenly on the injected leaves, and incubation was continued for 4 h. Total proteins were extracted from transiently expressed tobacco leaves using extraction buffer (50 mM Tris/HCl (pH 7.5), 2 mM MgCl_2_, 5 mM DTT, 150 mM NaCl, 20% glycerol, 0.1% Nonidet P-40, 100 uM protease inhibitor cocktail), and immunoblotting using anti-GFP (Abmart, Shanghai, China), anti-Flag antibodies (Abmart, Shanghai, China) and anti-His antibodies (Abmart, Shanghai, China) were used for further analysis.

## Acknowledgments

This study was supported by the key projects at the central government level: The Ability Establishment of Sustainable Use for Valuable Chinese Medicine Resources (2060302), the R & D project of ’ Jianbing ’ and ’ Lingyan ’ in Zhejiang Province (2022C02023), and National Natural Science Foundation of China (81773835).

## Author contributions

SL performed the experiments, analyzed the data, and wrote the first draft. JL and AA modified language. XY Performed some experiments. ZL and JD designed experiments and edited the manuscript.

